# *Erbb4* deletion from fast-spiking interneurons causes psychosis-relevant neuroimaging phenotypes

**DOI:** 10.1101/2022.03.07.483347

**Authors:** A. Kiemes, M. E. Serrano Navacerrada, E. Kim, K. Randall, C. Simmons, L. Rojo Gonzalez, M.M. Petrinovic, D.J. Lythgoe, D. Rotaru, D. Di Censo, L. Hirshler, E. L. Barbier, A. C. Vernon, J. M. Stone, C. Davies, D. Cash, G. Modinos

## Abstract

Converging lines of evidence suggest that dysfunction of cortical parvalbumin-expressing (PV+) GABAergic interneurons is a core feature of psychosis. This dysfunction is thought to underlie neuroimaging abnormalities commonly found in patients with psychosis, particularly in the hippocampus. These include increases in resting cerebral blood flow (CBF) and levels of glutamatergic metabolites, and decreases in binding of GABA_A_ α5 receptors and the synaptic density marker synaptic vesicle glycoprotein 2A (SV2A). However, direct links between PV+ interneuron dysfunction and these neuroimaging readouts have yet to be established. Conditional deletion of a schizophrenia susceptibility gene, the tyrosine kinase receptor *Erbb4*, from cortical and hippocampal PV+ interneurons leads to several synaptic, behavioral and cognitive phenotypes relevant to psychosis in mice. Here, we investigated how this PV+ interneuron disruption affects the hippocampal *in vivo* neuroimaging readouts in the *Erbb4* model. Adult *Erbb4* conditional mutant mice (*Lhx6-Cre;Erbb4*^*F/F*^, n=12) and their wild-type littermates (*Erbb4*^*F/F*^, n=12) were scanned in a 9.4T magnetic resonance scanner to quantify CBF and glutamatergic metabolite levels (glutamine, glutamate, GABA). Subsequently, we assessed GABA_A_ receptors and SV2A density using quantitative autoradiography. *Erbb4* mutant mice showed significantly elevated CBF and glutamine levels, as well as decreased SV2A density compared to wild-type littermates. No significant GABA_A_ receptor density differences were identified. These findings demonstrate that specific disruption of cortical PV+ interneurons in mice recapitulate some of the key neuroimaging findings in psychosis patients, and link PV+ interneuron deficits to non-invasive, translational measures of brain function and neurochemistry that can be used across species.

## Introduction

Multiple lines of evidence suggest that inhibitory GABAergic interneuron dysfunction is a core feature of psychosis^1^, and that this dysfunction underlies the abnormalities in hippocampal activity commonly observed in the disorder^2^. More specifically, *post-mortem* human brain studies in psychosis have identified reductions in the GABA-synthesizing enzyme GAD67^3^, inhibitory interneuron number^4^, calcium-binding protein parvalbumin (PV) expressed by some GABAergic interneurons^5,6^, and increases in GABA_A_ receptor density^7^. Among these, PV+ interneurons, a type of fast-spiking GABAergic cells that modulate neural network oscillations at the gamma frequency^8^, have been implicated in the pathophysiology of psychosis^9–11^. Abnormal gamma oscillations have been identified in individuals with psychosis spectrum disorders^12,13^ and are thought to underlie their cognitive symptoms^14^. Experiments in a neurodevelopmental animal model (methylazoxymethanol model, MAM) demonstrated that PV+ interneuron loss in the hippocampus leads to psychosis-relevant neurophysiological and cognitive deficits (i.e. reduced oscillatory activity and impaired latent inhibition)^11^. These findings led to the hypothesis that PV+ interneuron dysfunction in the hippocampus plays a critical role in the pathophysiology of psychosis^15,16^. Briefly, PV+ disruption in the hippocampus disinhibits glutamatergic excitatory cell activity, resulting in local hyperactivity. This drives an increase in striatal dopamine through descending projections, proposed to underlie psychosis symptoms. A hyperactive and dysrhythmic hippocampus can also interfere with the function of hippocampal-prefrontal cortex projections, disrupting prefrontal activity and rhythmicity, leading to cognitive deficits^15,16^.

In humans, neuroimaging studies have identified hippocampal abnormalities consistent with a fundamental role of GABAergic dysfunction in the pathophysiology of psychosis. Patients with psychosis exhibit hippocampal hyperactivity as indexed by increased regional cerebral blood flow (CBF)^17,18^ and cerebral blood volume (CBV)^19–23^. Such hyperactivity has been linked to higher severity of positive symptoms such as delusions and hallucinations^2,15,24^. Increases in CBF are also observed in psychosis vulnerability states, including individuals at clinical high-risk (CHR) for psychosis and healthy individuals with high schizotypy^25–28^. As mentioned above, such activity increases are proposed to result from GABAergic interneuron dysfunction^2^. Supporting this premise, a positron emission tomography (PET) study with the non-selective GABA_A_ receptor (α1-3;5GABA_A_R) tracer [^11^C]flumazenil found increases in *in vivo* GABA_A_ receptor binding in antipsychotic-naïve psychosis patients, that were linked to their cognitive symptoms and abnormal cortical oscillations^29^. More recently, studies using the more selective PET radiotracer [^11^C]Ro15-4513, reported binding decreases in hippocampal GABAA α5 receptors (α5GABA_A_R) in antipsychotic-free patients^30^ but not in patients currently undergoing treatment^30,31^. Seeking to further characterize the nature of hippocampal dysfunction in psychosis, reductions in the synaptic vesicle glycoprotein 2A (SV2A) – a putative marker of synaptic density – have been reported in the hippocampus of patients by *in vivo* [^11^C]UCB-J PET imaging^32,33^. This corroborates post-mortem^34–39^ findings of decreased dendritic spines and synaptic markers, as well as genetic evidence of variants in synaptic protein coding genes^40–43^. Finally, other studies using proton magnetic resonance spectroscopy (^1^H-MRS) to quantify excitatory and inhibitory metabolites in the hippocampus identified increases in the levels of combined glutamine and glutamate (Glx)^44,45^, but not GABA^46,47^, in patients with psychosis compared to healthy controls. Despite these recent human neuroimaging advances supporting a key mechanistic role for GABAergic dysfunction in psychosis, such neuroimaging assessments cannot inform whether these signal changes are associated with specific neuronal subpopulations, such as PV+ interneurons.

One way to address this issue is by targeted (e.g. genetic) modification of specific cell types in animal models. This allows the effects of such genetic modifications to be assessed using the same neuroimaging modalities used in human (clinical) studies^48,49^, providing more direct evidence linking cellular defects to macroscopic *in vivo* neuroimaging changes. For example, previous work in the cyclin D2 knockout model identified increased CBV as a result of hippocampal PV+ interneuron reduction^50^. In another mouse model, deletion of the tyrosine kinase receptor *Erbb4* (a susceptibility gene linked to psychosis^51,52^) from PV+ interneurons^53,54^ in the cortex and hippocampus leads to several psychosis-relevant phenotypes^55–58^. These include synaptic deficits (e.g., decreased interneuron signaling in the hippocampus, dysregulated glutamatergic activity of hippocampal pyramidal cells)^55^, elevated striatal dopamine^58^ and psychosis-relevant behaviors (e.g., hyperlocomotion, impaired prepulse inhibition, impaired cognitive and social behavior)^55^. *Erbb4* mutant mice thus represent a suitable model in which to analyze the contribution of PV+ interneuron dysfunction to hippocampal abnormalities associated with psychosis using non-invasive, clinically translational neuroimaging methods.

Our study used the *Erbb4* mouse model to determine how the PV+ interneuron dysfunction affects *in vivo* functional neuroimaging readouts commonly used in humans: arterial spin labeling (ASL) to measure CBF, and ^1^H-MRS to measure glutamate, glutamine and GABA levels in the hippocampus. Next, we sought to characterize hippocampal receptor and synaptic densities in this model, using *ex vivo* quantitative autoradiography with radioligands previously used in human *in vivo* PET studies: [^3^H]Ro15-4513 to measure α5GABA_A_R, [^3^H]flumazenil for α1-3;5GABA_A_R, and [^3^H]UCB-J for SV2A. Based on the synaptic deficits previously reported in these animals^55^, and the evidence that PV+ interneuron deficits underlie hippocampal hyperactivity in psychosis^2^, we hypothesized that *Erbb4* mouse mutants would show increases in CBF, glutamatergic metabolites and α1-3;5GABA_A_R density, as well as decreases in α5GABA_A_R and SV2A density in the hippocampus compared to wild-type littermate controls.

## Methods

### Animals

All animal procedures were performed in accordance with UK Home Office Animals (Scientific Procedures) Act 1986 and approved by the local King’s College London Animal Welfare Ethical Review Body. Animals were maintained under standard laboratory conditions on a 12:12 light/dark cycle with water and food *ad libitum*. Mice carrying loxP-flanked *Erbb4* alleles^56^ were crossed with *Lhx6-Cre* mice^59^ to generate *Lhx6-Cre;Erbb4*^*F/F*^ conditional mutants. Wild-type *Erbb4*^*F/F*^ littermates were used as controls.

### Experimental design

Twelve *Lhx6-Cre;ErbB4*^*F/F*^ (9 female; 3 male) and 12 *Erbb4*^*F/F*^ control (5 female; 7 male) adult (PD98 ± 11 days) mice underwent *in vivo* MR imaging. MR images were acquired using a 9.4T Bruker BioSpec 94/20 scanner with an 86-mm volume transmission coil and receive-only 2×2 surface array coil. All MR data were acquired from anesthetized animals (see Anesthesia section below) in a single scanning session and the brains were collected immediately after scanning for quantitative autoradiography.

### Anesthesia

Mice were initially anesthetized with 5% isoflurane in a mixture of 70% air and 30% oxygen. Once they were positioned on the scanner bed, a subcutaneous bolus of medetomidine (0.05 mg/kg) was administered and the isoflurane reduced to 1.5%. Eight minutes after the bolus, a subcutaneous infusion of medetomidine (0.1 mg/kg/hr) was started and maintained until the end of the ASL scan^60,61^. Then, the medetomidine infusion was stopped and the isoflurane level was increased to 2% for the remaining scans.

### Arterial Spin Labeling

Pseudo-continuous ASL (pCASL) was used to quantify CBF. The pCASL protocol includes a perfusion scan and two pre-scans to determine the optimal label and control phase increments and an inversion efficiency (IE) scan for each mouse^62^. The labeling slice was positioned 5 mm upstream of the carotid bifurcation identified by localizer scans of the neck. The labeling duration (*τ*) and post-label delay were 3000/300 ms, 1500/300 ms, and 200/0 ms for the perfusion scan, pre-scans, and IE scan, respectively. The pre-scans and perfusion scan used a 2D spin-echo echo-planar imaging readout: echo time (TE)/repetition time (TR) = 14.1/4000 ms, readout bandwidth = 300 kHz, matrix = 92×60, field-of-view (FOV) = 18.4×12 mm. Ten 1-mm-thick slices were acquired for the perfusion scan, and a single 4-mm-thick slice for the pre-scans. For the IE scan, a single 1-mm-thick slice 3 mm downstream of the labeling slice was acquired using a flow-compensated gradient echo sequence: TE/TR = 5.2/220 ms, flip angle (FA) = 25°, matrix = 200×180, FOV = 20×18 mm, 4 averages. The perfusion scan comprised 40 label/control image pairs. Four additional control images were acquired with reversed phase-encoding blips for distortion correction, which was performed using FSL topup (v5.0.10^63^).

T1 maps were acquired for CBF quantification using an MP2RAGE sequence: TE/TR = 2.5/7 ms, MP2RAGE_TR_ = 7000 ms, inversion times TI1/TI2 = 800/2500 ms, FA = 7/7°, matrix = 108×108×64, FOV = 16.2×16.2×9.6 mm. The qi_mp2rage command from the QUantitative Imaging Toolbox (QUIT v2.0.2^64^) was used to compute T1 maps from the complex MP2RAGE images.

Custom MATLAB scripts were written to calculate the mean IE in manually drawn regions of interest (ROIs) around both carotid arteries and quantitative CBF maps using the following equations:

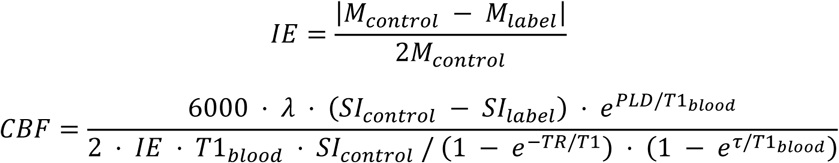

M_control_ and M_label_ are the complex signals from the control and label images from the IE scan, SI_control_ and SI_label_ are the time-averaged signal intensities of the control and label images from the perfusion scan, assuming the blood-brain partition coefficient *λ*= 0.9 ml/g, and T1_blood_ = 2.4 s.

The T1 images were used to register all subjects to the Allen mouse brain Common Coordinate Framework v3 (CCFv3) using antsRegistration to perform sequential rigid-body, affine, and SyN diffeomorphic registrations (ANTs v2.1.0^65^). CBF maps were normalized by the mean CBF of the whole brain, and then mean CBF values were calculated for 21 ROIs derived from the CCFv3 atlas labels. We focused our analyses on the dorsal and ventral hippocampus (Figure 2A). For completeness, exploratory independent t-tests of other atlas-derived ROIs are presented in the supplementary materials (Table S2).

### Magnetic resonance spectroscopy

^1^H-MRS was used to quantify hippocampal metabolite profiles^66^ in conditional *Erbb4* mouse mutants and control animals. After manually placing the voxel at the hippocampus (Figure 3A), with the aid of T1 structural images, individual spectra were acquired using a Point REsolved Spectroscopy (PRESS) pulse sequence^67^ with the following parameters: TE = 8.23 ms, TR = 2500 ms, 512 averages, acquisition bandwidth = 4401 Hz, 2048 acquisition points, voxel size = 1.5×1.5×3 mm. Outer volume suppression and water suppression with variable pulse power and optimized relaxation delays (VAPOR) were used in order to mitigate the contribution of signal from outside the prescribed voxel and suppress unwanted signal from water.

MR spectra were analyzed with two software packages: FID Appliance (FID-A^68^) and Linear Combination (LC) Model version 6.3^69,70^. First, FID-A was used to pre-process ^1^H-MRS data, simulate the metabolites and create a basis set (model spectra). Then, we used LCModel to calculate the water-referenced concentration (in mM) of the different metabolites by applying linear combinations of the model spectra to determine the best fit of the individual ^1^H-MRS data^71^. Finally, the method of Cramér Rao (Cramér Rao Lower Bound, CRLB) was applied to ensure the reliability of the metabolite quantification, by which metabolite concentrations with S.D. ≥ 20% are classified as not accurately detectable and are discarded^72,73^. Using these criteria no data had to be discarded for our metabolites of interest: gamma-aminobutyric acid (GABA), glutamine (Gln), and glutamate (Glu) (Figure 1).

**Figure 1.**
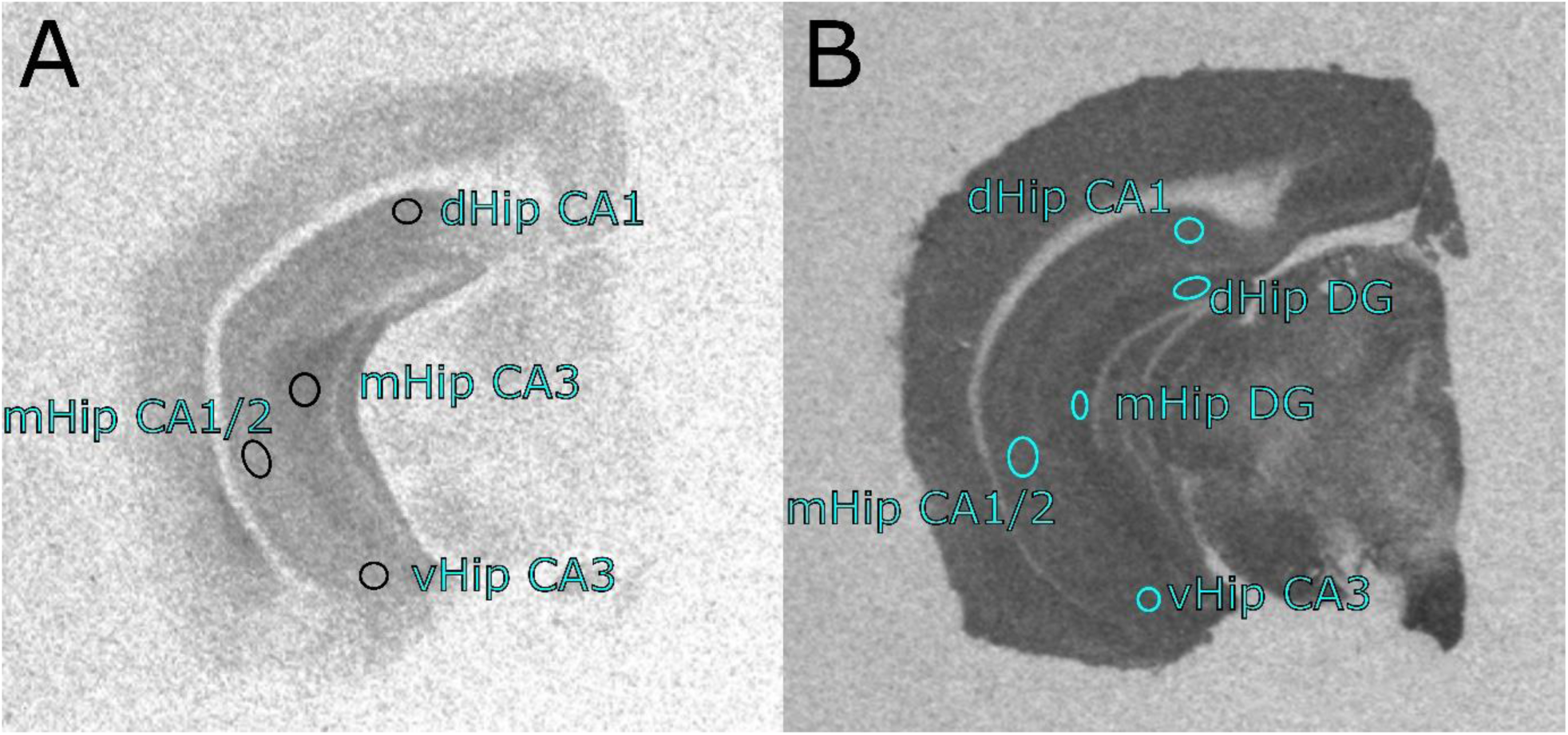
Representative hippocampal regions of interests. (A) Regions of interests sampled for [^3^H]Ro15-4513 and [^3^H]flumazenil. (B) Regions of interests sampled for [^3^H]UCB-J. dHip CA1: dorsal hippocampus CA1; mHip CA3: middle hippocampus CA3; mHip CA1/2: middle hippocampus CA1/CA2; vHip CA3: ventral hippocampus CA3; dHip DG: dorsal hippocampus dentate gyrus; mHip DG: middle hippocampus dentate gyrus.

### Quantitative autoradiography

Following the MR scanning, mice were sacrificed, the brains dissected, and flash frozen in cold (−40°C) isopentane on dry ice, then stored at -80°C. Frozen brains were coronally cryosectioned at 20µm and mounted onto glass slides, then dried on a hotplate. Quantitative autoradiography was performed as previously described^74,75^ using radioligands [^3^H]Ro15-4513, [^3^H]flumazenil and [^3^H]UCB-J. All slides were soaked in Tris buffer (50mM) for 20 minutes prior to incubation with radioligands for specific or non-specific binding, and this was followed by two washes in Tris buffer for 2 minutes each, and a rinse in dH_2_O, before overnight air-drying.

To quantify density of α5GABA_A_R^76–78^ sections were incubated for 60 minutes at room temperature in 2nM [^3^H]Ro15-4513 (Perkin Elmer, NET925250UC), or in 2nM [^3^H]Ro15-4513 with 10 µM bretazenil (Sigma, B6434) for nonspecific binding. To quantify α1-3;5GABA_A_R^79^ sections were incubated for 60 minutes at 4°C in 1nM [^3^H]flumazenil (Perkin Elmer, NET757001MC), or in 1nM [^3^H]flumazenil with 10 µM flunitrazepam (Sigma Aldrich, F-907 1ML) for nonspecific binding. To quantify SV2A density^80^, sections were incubated for 60 minutes at room temperature in 3nM [^3^H]UCB-J (Novandi Chemistry AB, NT1099), or in 3nM [^3^H]UCB-J with 1mM levetiracetam (Sigma Aldrich, L8668) for nonspecific binding.

Dried slides and [^3^H] standards (American Radiolabelled Chemicals, Inc., USA, ART-123A) were placed into light-proof cassettes, and a [^3^H]-sensitive film (Amersham Hyperfilm, 28906845) was placed on top. The films were exposed 2 weeks for [^3^H]UCB-J, 4 weeks for [^3^H]flumazenil and 8 weeks for [^3^H]Ro15-4513. All films were developed with a Optimax 2010 film developer (Protec GmbH & Co, Germany) and autoradiographs captured using an AF-S Micro NIKKOR 60mm lens on top of a light box (Northern Lights, USA). Lighting conditions were kept the same during imaging capture of each film. Optical density was measured in standards and ROIs of autoradiographs using ImageJ (1.52e). Receptor binding (µCi/mg) was calculated with robust regression interpolation in GraphPad Prism (v9.2.0 for Windows) using standard curves created from optical density measurements of [^3^H]-standards slide for each film.

For [^3^H]Ro15-4513 and [^3^H]flumazenil, four ROIs were sampled: CA1 of the dorsal hippocampus, CA3 of the middle hippocampus, CA1/2 of the middle hippocampus, and the CA3 of the ventral hippocampus (Figure 1A). Owing to better signal/contrast to noise ratio of [^3^H]UCB-J autoradiographs (Figure 1B), we were also able to analyze the binding in the dentate gyrus. These hippocampal ROIs were selected based on previous evidence implicating their involvement in psychosis^19,22,24,55,81,82^. For completeness, further non-hippocampal ROIs (amygdala, retrosplenial cortex, visual cortex, prelimbic cortex, motor cortex, orbital cortex) were sampled, and their exploratory statistical analysis for all three radioligands is presented in the supplementary materials (Table S3-5).

### Statistical Analysis

Statistical analysis was conducted using GraphPad Prism software (v9.2.0 for Windows). To investigate the group differences in CBF and autoradiography data, we used mixed-effects analyses, with the genotype (*Lhx6-Cre;Erbb4*^*F/F*^ vs *Erbb4*^*F/F*^ control mice) as between-group factor and ROI as within-group factor. Any significant interaction effects of genotype x ROI were investigated by follow-up multiple comparisons and *p* values were adjusted using Bonferroni correction (*p*_*corr*_). For metabolite data, we analyzed group differences using independent t-tests per metabolite and Bonferroni-adjusted *p* values. Significance threshold was set to *p* < 0.05. Cohen’s d and partial eta squared effect sizes were calculated from test statistics using the *effectsize* library (v 0.5^83^) in RStudio (v1.3.1093). Due to technical failures such as scanning faults, inadequate tissue preparation, and Covid-19 restrictions limiting laboratory access, the following mouse data were missing: 2 *Erbb4*^*F/F*^ mice for CBF, 3 *Erbb4*^*F/F*^ and 1 *Lhx6-Cre;Erbb4*^*F/F*^ mice for [^3^H]UCB-J autoradiography, 1 *Erbb4*^*F/F*^ and 2 *Lhx6-Cre;Erbb4*^*F/F*^ mice for [^3^H]-Ro15-4513, and 2 *Erbb4*^*F/F*^ and 3 *Lhx6-Cre;Erbb4*^*F/F*^ mice for [^3^H]flumazenil. To better graphically depict the comparison between different ^1^H-MRS metabolites (Figure 3B), we calculated *z* scores of individual concentrations in relation to the pooled group mean metabolite concentration.

## Results

### Ventral hippocampal CBF is increased in *Lhx6-Cre;Erbb4*^*F/F*^ mice

There was no significant main effect of genotype on CBF (*F*_(1,20)_=2.75, *p*=0.11,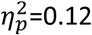), but we identified a genotype × ROI interaction effect (*F*_(1,20)_=11.91, *p*=0.003, 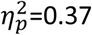). Follow-up analysis (Figure 2B) revealed a significant increase in CBF values in *Lhx6-Cre;Erbb4*^*F/F*^ mice compared to wild-type littermates in the ventral (*t*_(40)_=2.54, *p*_*corr*_=0.03, *d*=0.80), but not in the dorsal hippocampus (*t*_(40)_=0.67, *p*_*corr*_>0.9, *d*=0.21).

**Figure 2.**
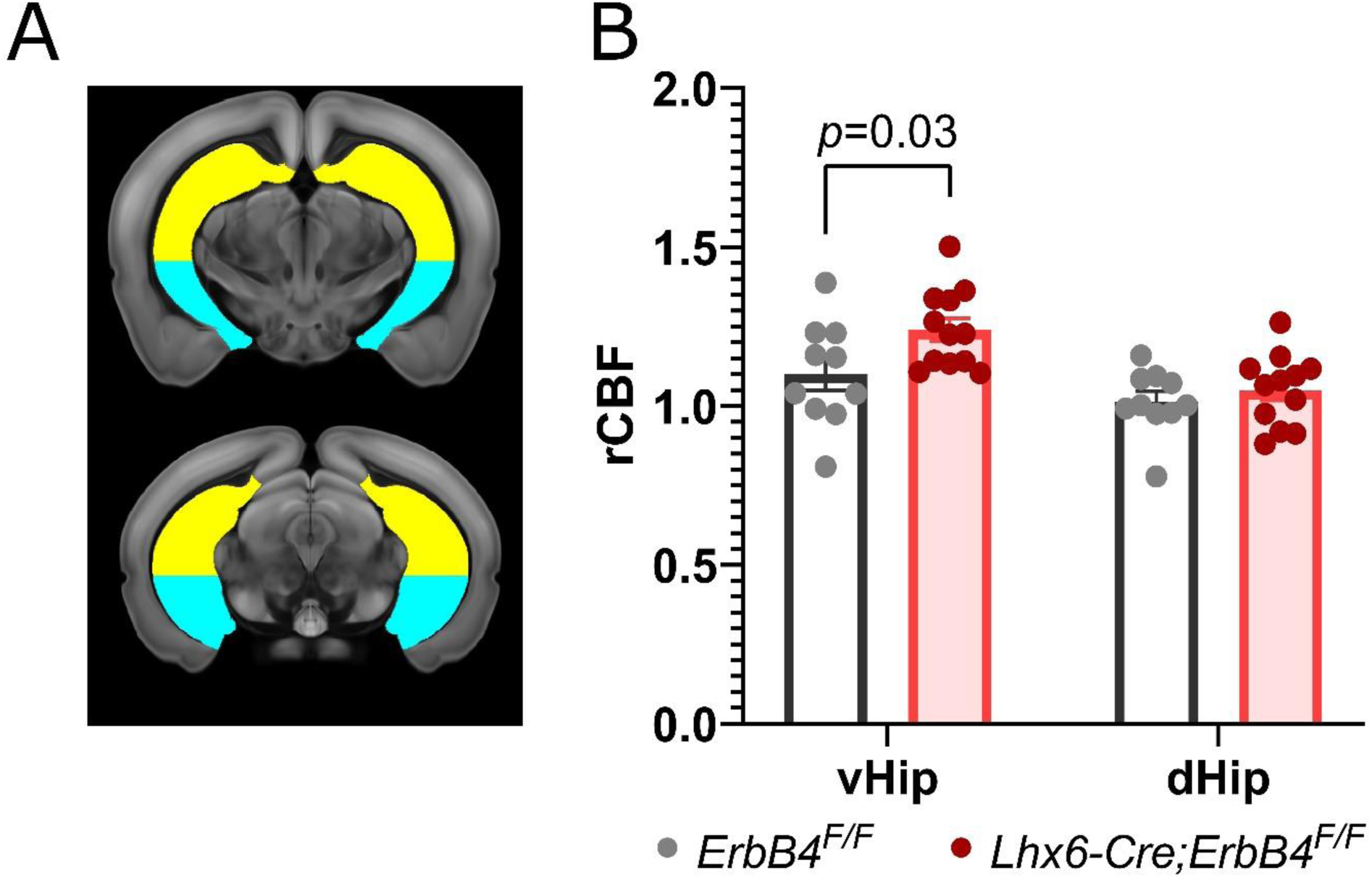
Regional cerebral blood flow in Lhx6-Cre;Erbb4^F/F^ mice is increased in the ventral hippocampus. (A) Hippocampal regions of interest for sampling overlaid on the mouse brain template (approximate distance from Bregma^84^, top -2.8, bottom -3.2); yellow = dorsal hippocampus and blue = ventral hippocampus. (B) Regional cerebral blood flow is increased in the ventral hippocampus, but not in the dorsal hippocampus in Lhx6-Cre;Erbb4^F/F^ mutants (n=12) compared to control mice (n=10; multiple comparison independent t-tests). vHip: ventral hippocampus; dHip: dorsal hippocampus.

### Glutamine levels are increased in ventral hippocampus of *Lhx6-Cre;Erbb4*^*F/F*^ mice

*Lhx6-Cre;Erbb4*^*F/F*^ mice showed increased glutamine levels in the ventral hippocampus compared to control animals (*t*_(22)_=4.60, *p*<0.001, *d*=1.96, Figure 3B and Table 1). There were no significant group differences in either glutamate or GABA concentrations (Table 1).

**Figure 3.**
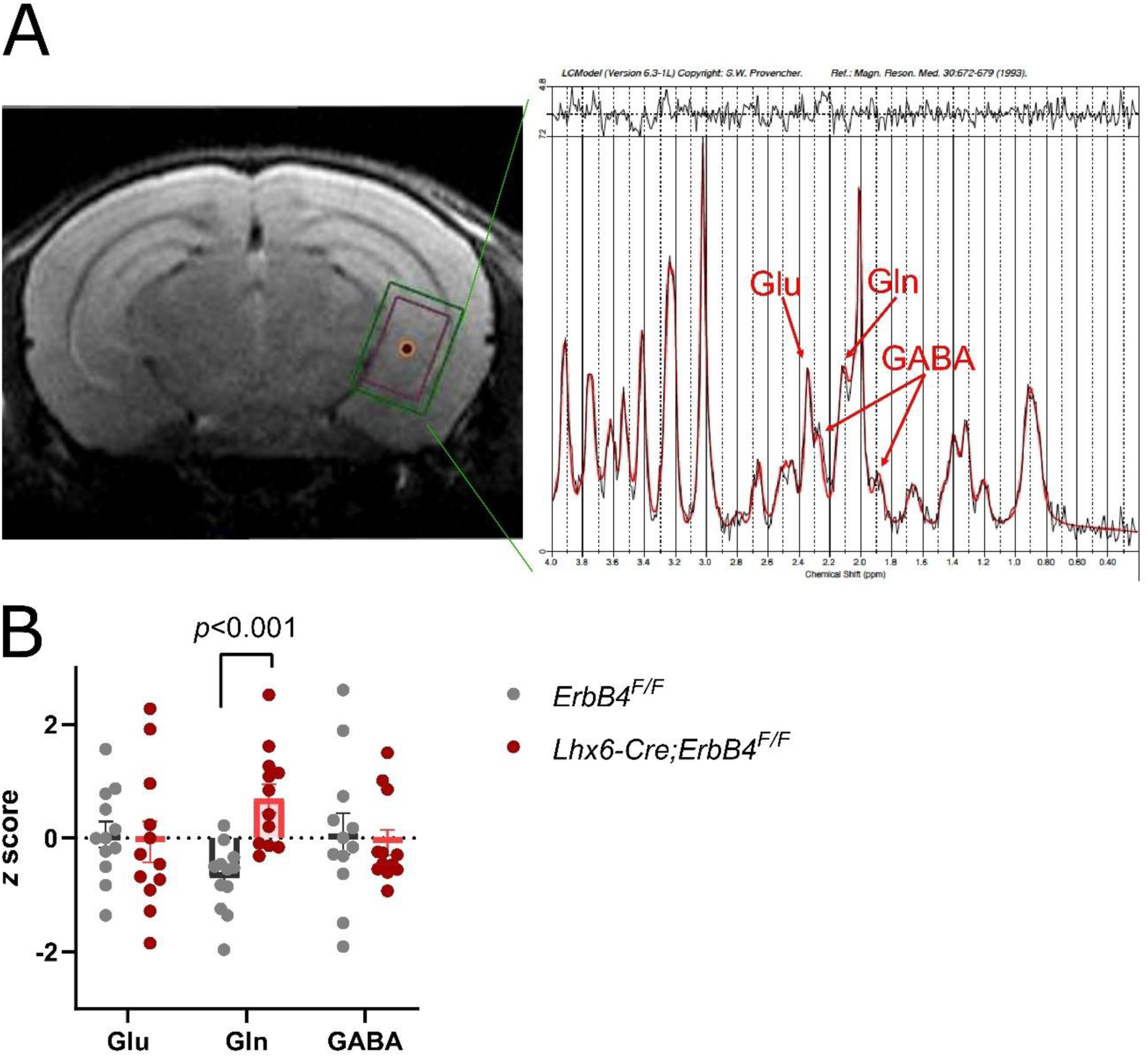
(A) Representative ^1^H-MRS PRESS voxel (red) and corresponding shim (green) placement in ventral hippocampus (left) and ^1^H-MRS spectrum (right). (B) Z-scores of ^1^H-MRS metabolites in the ventral hippocampus. Glutamine was significantly increased in Lhx6-Cre;Erbb4^F/F^ mutant mice (n=12) compared to control mice (n=12; independent t-tests). GABA: gamma-aminobutyric acid; Glu: glutmate; Gln: glutamine.

**Table 1.** ^1^H-MRS absolute metabolite concentrations in millimolar

|  | *ErbB4<sup>F/F</sup>*<br>(n=12) | *Lhx6-Cre;ErbB4<sup>F/F</sup>*<br>(n=12) | *ErbB4<sup>F/F</sup>* vs. *Lhx6-Cre;ErbB4<sup>F/F</sup>* |  |
| --- | --- | --- | --- | --- |
|  | Mean (SD) | Mean (SD) | *t* | *p* * |
| Glu | 6.77 (0.51) | 6.68 (0.80) | 0.31 | >0.9 |
| Gln | 2.29 (0.23) | 2.84 (0.34) | 4.60 | <0.001 |
| GABA | 2.08 (0.49) | 2.02 (0.30) | 0.38 | >0.9 |

Glu: glutamate; Gln: glutamine; \*Bonferroni-adjusted *p* values

### *Lhx6-Cre;Erbb4*^*F/F*^ mice display decreased [^3^H]-UCB-J binding in the hippocampus

There was a significant main effect of genotype on [^3^H]UCB-J binding (F_(1,18)_=7.27, *p*=0.02, 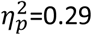), indicating reduced synaptic density in *Lhx6-Cre;Erbb4*^*F/F*^ mice compared to control animals across all hippocampal ROIs (Figure 4A). No genotype x ROI interaction effect was observed (F_(4,66)_=0.68, *p*=0.61, 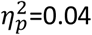).

**Figure 4.**
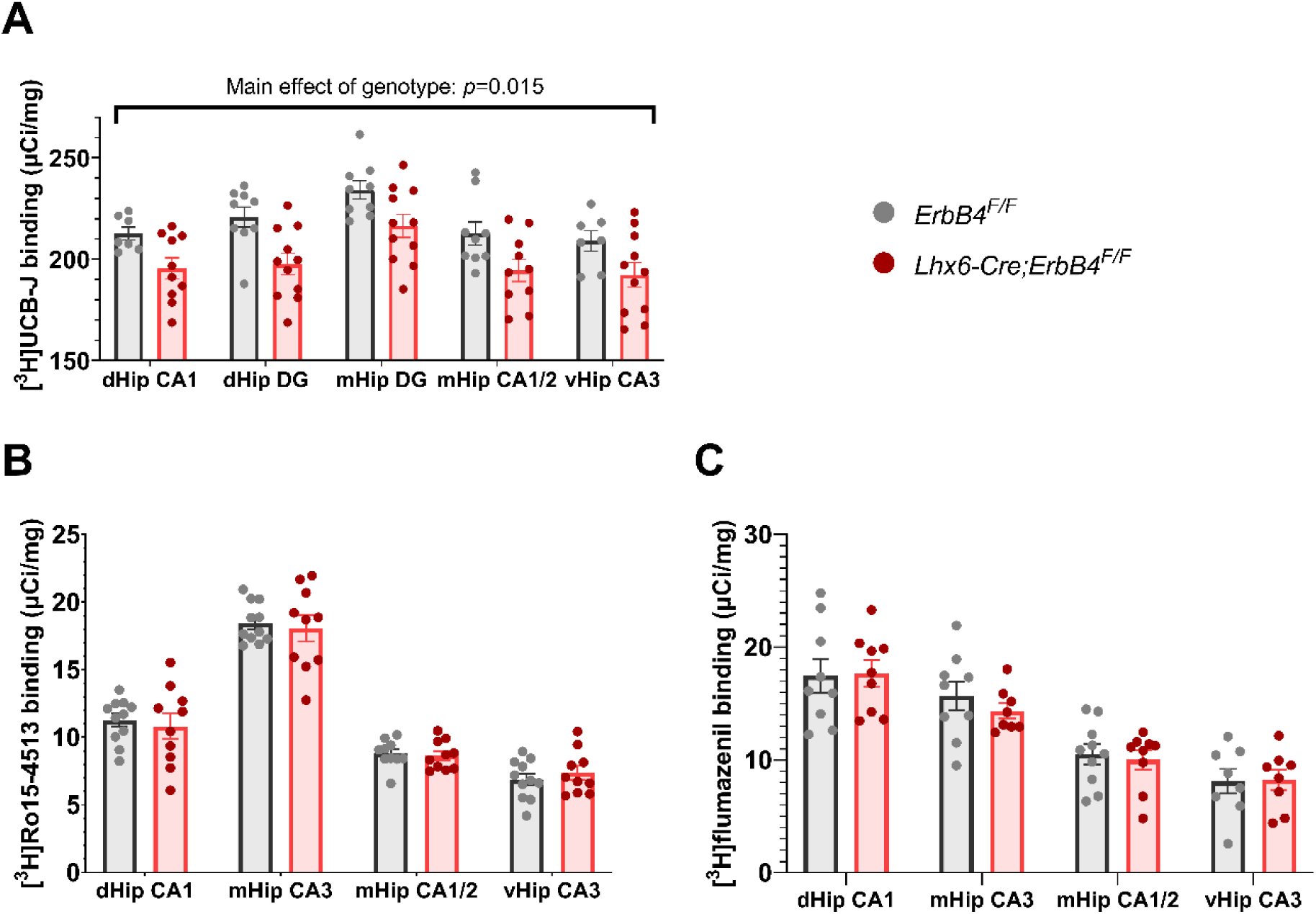
[^3^H]UCB-J, but not [^3^H]Ro15-4513 or [^3^H]flumazenil binding in Lhx6-Cre;Erbb4^F/F^ mice is decreased across the hippocampus. (A) [^3^H]UCB-J showed a significant decrease in synaptic density in Lhx6-Cre;Erbb4^F/F^ mice (n=11) compared to control animals (n=9) across all hippocampal ROIs (mixed effects analysis; main effect of genotype: F_(1,18)_=7.27, *p*=0.02,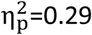). (B) [^3^H]Ro15-4513 (n=10-11/group) and (C) [^3^H]flumazenil (n=9-10/group) binding did not differ by genotype. dHip CA1: dorsal hippocampus CA1; mHip CA3: middle hippocampus CA3; mHip CA1/2: middle hippocampus CA1/CA2; vHip CA3: ventral hippocampus CA3; dHip DG: dorsal hippocampus dentate gyrus; mHip DG: dentate gyrus.

[^3^H]Ro15-4513 binding, as a measure of α5GABA_A_R density, did not differ significantly between the two genotypes (F_(1,19)_=0.05, *p*=0.82,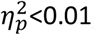 Figure 4B). Similarly, α1-3;5GABA_A_R density as measured by [^3^H]flumazenil binding did not differ between the genotypes (F_(1,17)_=0.07, *p*=0.79, 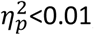Figure 4C).

## Discussion

In this study, we used conditional *Erbb4* mutants to examine the effects of PV+ inhibitory interneuron dysfunction on key neuroimaging markers of psychosis. We identified abnormalities in hippocampal activity, neurochemistry and synaptic density that are largely convergent with clinical neuroimaging findings in patients. More specifically, compared to wild-type mice, *Erbb4* mutants showed increased CBF and glutamine levels in the ventral hippocampus, as well as decreases in SV2A levels across hippocampal sub-regions. GABA and glutamate levels did not significantly differ between the groups, and there were no differences in the density of any of the measured GABAergic receptors. Interestingly, in our exploratory analysis of regions outside of the hippocampus (see Supplementary Tables S1, S3-5 and Supplementary Discussion) we found additional group differences, although these did not survive multiple comparisons correction. Thus, in *Erbb4* mutant mice, we observed further elevations of CBF in the entorhinal cortex and increases of α5GABA_A_R density in the retrosplenial cortex. Additionally, decreases in α1-3;5GABA_A_R density in the retrosplenial cortex approached significance.

Our investigation focused primarily on the hippocampus, based on pre-existing hypotheses suggesting that PV+ interneuron loss in the ventral part of this region contributes to its hyperactivity and is associated with further electrophysiological and cognitive deficits relevant to psychosis^2,15^. Indeed, hippocampal disinhibition is suggested to disrupt cognitive functioning in schizophrenia^15^, consistent with the vital role of PV+ interneurons in entraining gamma oscillations^8^. In concordance with the hippocampal hyperactivity hypothesis^2^, in the *Erbb4* model of PV+ interneuron dysfunction, inhibitory control over pyramidal neurons is disrupted^55^, leading to increased neural activity captured by CBF via neurovascular coupling^85^. Importantly, our findings are localized to the ventral part of the hippocampus, matching the previous findings of increased CBV in psychosis patients^19,21–23^ and CBF in CHR patients^26,28^ in the human anatomical equivalent, the anterior hippocampus. Further, our findings align with previous evidence of increased CBV in the cyclin D2 knockout model that exhibits PV+ interneuron loss^50^. Here, we used a model that manipulates an established schizophrenia susceptibility gene instead, and expand findings of PV+ interneuron related hippocampal hyperactivity in the less invasive and quantitative MR method of ASL.

In terms of our ^1^H-MRS findings, we identified an increase in glutamine, but not glutamate or GABA, in the ventral hippocampal region. Disinhibition of pyramidal neuronal activity in the *Erbb4* model^53,55,86^, is thought to lead to increased glutamate release^87,88^. However, glutamine, a precursor of glutamate, could be considered a better indicator of glutamatergic neurotransmission^89^. This is based on the premise that any synaptically released glutamate is quickly taken up by astrocytes and recycled to glutamine^90,91^. Accordingly, increased glutamine in the medial temporal lobe / hippocampi has previously been detected by ^1^H-MRS in psychosis patients^44^. Other human studies also showed evidence of elevated Glx^44,45^ -a composite of glutamate and glutamine – that is preferentially measured at lower magnetic fields such as 1.5 or 3T in humans, where the separation between those two metabolites is not robust^92^. These findings suggest that increased glutamine may be a good indicator of elevated glutamatergic neurotransmission and it occurs as a consequence of PV+ interneuron dysfunction.

Furthermore, no *Erbb4* genotype effect was observed in hippocampal ^1^H-MRS GABA levels. Previous findings in conditional *Erbb4* mutants had identified reduced expression of two GABA synthesizing GAD isomers, GAD65 and GAD67, as well as reduced frequency of miniature inhibitory postsynaptic GABAergic currents^55^. However, it is also known that, as a result of PV+ inhibitory neuron disruption, both PV+ interneurons and excitatory pyramidal cells eventually become hyperactive in *Erbb4* mutants through possible compensatory mechanism in order to maintain excitation/inhibition balance^55^. This compensatory inhibitory interneuron activity may counteract any deficits in GABA synthesis, thereby explaining the lack of measurable differences in GABA between the groups. Indeed, no changes in hippocampal GABA levels were identified in psychosis patients by a previous ^1^H-MRS GABA study^46^.

Although we hypothesized changes in hippocampal GABAergic receptor densities as a result of PV+ interneuron disruption, we found no differences between *Erbb4* mutants and control mice in either α1-3;5GABA_A_R or α5GABA_A_R. In humans, increases in α1-3;5GABA_A_R availability^29^ and decreases in the more specific α5GABA_A_R^30^ have been identified in groups of antipsychotic-naïve and medication-free schizophrenia patients, respectively. As *Erbb4* deletion specifically affects PV+ interneurons, a lack of α5GABA_A_R changes may be due to this subunit’s putative co-expression with somatostatin-expressing rather than PV+ interneurons^78^, suggesting that perhaps PV+ interneurons are not responsible for the α5GABA_A_R changes seen in humans^30^. Contrastingly, imaging transcriptomics suggest that the distribution of flumazenil binding and PV+ interneuron expression are correlated^78^ thus we expected to see changes in α1-3;5GABA_A_R. However, while preclinical evidence in the *Erbb4* model has demonstrated small decreases in α1GABA_A_R clusters at PV+ interneuron terminals^55^, it is possible α1-3;5GABA_A_R are upregulated outside of such terminals as a compensatory mechanism due to decreased GABAergic neurotransmission^7,29^. Future studies should investigate α1-3;5GABA_A_R density in the *Erbb4* animal model longitudinally to understand whether compensatory increases develop as a result of PV+ interneuron dysfunction.

Finally, *post-mortem*^34–39^ and genetic^40–43^ evidence suggest that synaptic dysfunction plays an important role in psychosis pathophysiology. Recent clinical neuroimaging studies have provided *in vivo* evidence for synaptic density decreases in psychosis patients, using PET radioligand [^11^C]UBC-J to image the synaptic glycoprotein SV2A^32,33^, a putative marker of synaptic density. Synaptic deficits are present in *Erbb4* mutant mice: excitatory synapses onto fast-spiking inhibitory neurons and presynaptic boutons in chandelier cells^55^, which are highly expressed in regions such as the hippocampus^93^, are reduced. Our study shows that such synaptic losses can be measured at a macroscopic scale via autoradiography in rodents and suggest that PV+ interneuron dysfunction may be underlying the reductions of SV2A observed in patients with psychosis.

There are some limitations to our study. Despite known sex differences in psychosis such as incidence rate, age of illness onset, illness course and treatment response^94,95^, both male and female mice were used for our study. This was based on following best practice^96,97^ and the 3Rs^98^ to avoid sex bias in preclinical research^99^. Behavioral testing was not performed in our animals, which may have enabled investigating associations with the imaging data. However, the behavior of *Erbb4* mutant mice has already been well characterized^55–57,100^ and the scope of our study was limited to the neuroimaging phenotypes arising from PV+ interneuron dysfunction. Future studies may expand on these results and link neuroimaging with behavioral readouts to better understand their relationships in the context of this model system. Finally, the mice were imaged at only one time-point (adulthood). Future studies should capitalize on the repeatability of *in vivo* neuroimaging^49^ and inform developmental trajectories of PV+ interneuron dysfunction on neuroimaging phenotypes.

In summary, our study provides direct evidence linking PV+ interneuron dysfunction in the *Erbb4* mouse model to analogues of *in vivo* neuroimaging alterations previously identified in psychosis and CHR patients. These alterations include increased CBF and glutamine levels, as well as reduced synaptic density in the hippocampus. Overall, these findings suggest that the use of translational neuroimaging methods may be a viable strategy to identify new therapeutic targets and serve as non-invasive measures of target engagement. Furthermore, our findings support the view that targeting inhibitory dysfunction in the hippocampus may be a promising therapeutic strategy for psychosis.

## Acknowledgements

The authors would kindly like to thank Bernard Clemence and Beatriz Rico for kindly providing the animals in this study. We would also like to thank Beatriz Rico for generously contributing her expertise and input to the interpretation of findings.

## References

1. Marín O. Interneuron dysfunction in psychiatric disorders. Nat Rev Neurosci. 2012;13(2):107–120. doi:10.1038/nrn3155

2. Heckers S, Konradi C. GABAergic mechanisms of hippocampal hyperactivity in schizophrenia. Schizophrenia Research. 2015;167(1):4–11. doi:https://doi.org/10.1016/j.schres.2014.09.041

3. Heckers S, Stone D, Walsh J, Shick J, Koul P, Benes FM. Differential hippocampal expression of glutamic acid decarboxylase 65 and 67 messenger RNA in bipolar disorder and schizophrenia. Archives of general psychiatry. 2002;59(6):521–529. doi:10.1001/archpsyc.59.6.521

4. Benes FM, Kwok EW, Vincent SL, Todtenkopf MS. A reduction of nonpyramidal cells in sector CA2 of schizophrenics and manic depressives. Biological Psychiatry. 1998;44:88–97. doi:10.1016/S0006-3223(98)00138-3

5. Zhang ZJ, Reynolds GP. A selective decrease in the relative density of parvalbumin-immunoreactive neurons in the hippocampus in schizophrenia. Schizophrenia Research. 2002;55(1-2):1-10. doi:10.1016/S0920-9964(01)00188-8

6. Konradi C, Yang CK, Zimmerman EI, et al. Hippocampal interneurons are abnormal in schizophrenia. Schizophrenia Research. 2011;131(1-3):165-173. doi:10.1016/j.schres.2011.06.007

7. Benes FM, Khan Y, Vincent SL, Wickramasinghe R. Differences in the subregional and cellular distribution of GABAA receptor binding in the hippocampal formation of schizophrenic brain. Synapse. 1996;22(4):338–349. doi:10.1002/(SICI)1098-2396(199604)22:4<338::AID-SYN5>3.0.CO;2-C

8. Antonoudiou P, Tan YL, Kontou G, Upton AL, Mann EO. Parvalbumin and Somatostatin Interneurons Contribute to the Generation of Hippocampal Gamma Oscillations. J Neurosci. 2020;40(40):7668–7687. doi:10.1523/JNEUROSCI.0261-20.2020

9. Curley AA, Lewis DA. Cortical basket cell dysfunction in schizophrenia. J Physiol. 2012;590(4):715–724. doi:10.1113/jphysiol.2011.224659

10. Lewis DA. The chandelier neuron in schizophrenia. Dev Neurobiol. 2011;71(1):118–127. doi:10.1002/dneu.20825

11. Lodge DJ, Behrens MM, Grace AA. A loss of parvalbumin-containing interneurons is associated with diminished oscillatory activity in an animal model of schizophrenia. Journal of Neuroscience. 2009;29(8):2344–2354. doi:10.1523/JNEUROSCI.5419-08.2009

12. Sun Y, Farzan F, Barr MS, et al. Gamma oscillations in schizophrenia: Mechanisms and clinical significance. Brain research. 2011;1413:98–114. doi:https://doi.org/10.1016/j.brainres.2011.06.065

13. Reilly TJ, Nottage JF, Studerus E, et al. Gamma band oscillations in the early phase of psychosis: A systematic review. Neuroscience & Biobehavioral Reviews. 2018;90:381–399. doi:https://doi.org/10.1016/j.neubiorev.2018.04.006

14. Uhlhaas PJ, Singer W. Abnormal neural oscillations and synchrony in schizophrenia. Nat Rev Neurosci. 2010;11(2):100–113. doi:10.1038/nrn2774

15. Grace AA, Gomes FV. The Circuitry of Dopamine System Regulation and its Disruption in Schizophrenia: Insights Into Treatment and Prevention. Schizophrenia Bulletin. 2019;45(1):148–157. doi:10.1093/schbul/sbx199

16. Grace AA. Dysregulation of the dopamine system in the pathophysiology of schizophrenia and depression. Nature Reviews Neuroscience. 2016;17(8):524–532. doi:10.1038/nrn.2016.57

17. Liddle PF, Friston KJ, Frith CD, Hirsch SR, Jones T, Frackowiak RSJ. Patterns of Cerebral Blood Flow in Schizophrenia. British Journal of Psychiatry. 1992;160(2):179–186. doi:10.1192/bjp.160.2.179

18. Medoff DR, Holcomb HH, Lahti AC, Tamminga CA. Probing the human hippocampus using rCBF: Contrasts in schizophrenia. Hippocampus. 2001;11(5):543–550. doi:https://doi.org/10.1002/hipo.1070

19. Talati P, Rane S, Kose S, et al. Increased hippocampal CA1 cerebral blood volume in schizophrenia. NeuroImage: Clinical. 2014;5:359–364. doi:https://doi.org/10.1016/j.nicl.2014.07.004

20. Talati P, Rane S, Skinner J, Gore J, Heckers S. Increased hippocampal blood volume and normal blood flow in schizophrenia. Psychiatry Res. 2015;232(3):219–225. doi:10.1016/j.pscychresns.2015.03.007

21. McHugo M, Talati P, Armstrong K, et al. Hyperactivity and Reduced Activation of Anterior Hippocampus in Early Psychosis. Am J Psychiatry. 2019;176(12):1030–1038. doi:10.1176/appi.ajp.2019.19020151

22. Schobel SA, Chaudhury NH, Khan UA, et al. Imaging patients with psychosis and a mouse model establishes a spreading pattern of hippocampal dysfunction and implicates glutamate as a driver. Neuron. 2013;78(1):81–93. doi:10.1016/j.neuron.2013.02.011

23. Schobel SA, Kelly MA, Corcoran CM, et al. Anterior hippocampal and orbitofrontal cortical structural brain abnormalities in association with cognitive deficits in schizophrenia. Schizophrenia Research. 2009;114(1-3):110–118.

24. Grace AA. Dopamine System Dysregulation by the Ventral Subiculum as the Common Pathophysiological Basis for Schizophrenia Psychosis, Psychostimulant Abuse, and Stress. Neurotoxicity Research. 2010;18(3):367–376. doi:10.1007/s12640-010-9154-6

25. Allen P, Azis M, Modinos G, et al. Increased Resting Hippocampal and Basal Ganglia Perfusion in People at Ultra High Risk for Psychosis: Replication in a Second Cohort. Schizophrenia Bulletin. 2017;44(6):1323–1331. doi:10.1093/schbul/sbx169

26. Modinos G, Egerton A, McMullen K, et al. Increased resting perfusion of the hippocampus in high positive schizotypy: A pseudocontinuous arterial spin labeling study. Hum Brain Mapp. 2018;39(10):4055–4064. doi:10.1002/hbm.24231

27. Modinos G, Richter A, Egerton A, et al. Interactions between hippocampal activity and striatal dopamine in people at clinical high risk for psychosis: relationship to adverse outcomes. Neuropsychopharmacology. 2021;46(8):1468–1474. doi:10.1038/s41386-021-01019-0

28. Allen P, Chaddock CA, Egerton A, et al. Resting hyperperfusion of the hippocampus, midbrain, and basal ganglia in people at high risk for psychosis. American Journal of Psychiatry. 2016;173(4):392–399. doi:10.1176/appi.ajp.2015.15040485

29. Frankle WG, Cho RY, Prasad KM, et al. In vivo measurement of GABA transmission in healthy subjects and schizophrenia patients. Am J Psychiatry. 2015;172(11):1148–1159. doi:10.1176/appi.ajp.2015.14081031

30. Marques TR, Ashok AH, Angelescu I, et al. GABA-A receptor differences in schizophrenia: a positron emission tomography study using [11C]Ro154513. Molecular Psychiatry. April 2020. doi:10.1038/s41380-020-0711-y

31. Asai Y, Takano A, Ito H, et al. GABAA/Benzodiazepine receptor binding in patients with schizophrenia using [11C]Ro15-4513, a radioligand with relatively high affinity for α5 subunit. Schizophrenia Research. 2008;99(1):333–340. doi:10.1016/j.schres.2007.10.014

32. Onwordi EC, Halff EF, Whitehurst T, et al. Synaptic density marker SV2A is reduced in schizophrenia patients and unaffected by antipsychotics in rats. Nature Communications. 2020;11(1):246. doi:10.1038/s41467-019-14122-0

33. Radhakrishnan R, Skosnik PD, Ranganathan M, et al. In vivo evidence of lower synaptic vesicle density in schizophrenia. Mol Psychiatry. June 2021:1–9. doi:10.1038/s41380-021-01184-0

34. Garey LJ, Ong WY, Patel TS, et al. Reduced dendritic spine density on cerebral cortical pyramidal neurons in schizophrenia. J Neurol Neurosurg Psychiatry. 1998;65(4):446–453. doi:10.1136/jnnp.65.4.446

35. Glantz LA, Lewis DA. Decreased dendritic spine density on prefrontal cortical pyramidal neurons in schizophrenia. Arch Gen Psychiatry. 2000;57(1):65–73. doi:10.1001/archpsyc.57.1.65

36. Davidsson P, Gottfries J, Bogdanovic N, et al. The synaptic-vesicle-specific proteins rab3a and synaptophysin are reduced in thalamus and related cortical brain regions in schizophrenic brains. Schizophr Res. 1999;40(1):23–29. doi:10.1016/s0920-9964(99)00037-7

37. Matosin N, Fernandez-Enright F, Lum JS, et al. Molecular evidence of synaptic pathology in the CA1 region in schizophrenia. NPJ Schizophr. 2016;2:16022. doi:10.1038/npjschz.2016.22

38. Halim ND, Weickert CS, McClintock BW, et al. Presynaptic proteins in the prefrontal cortex of patients with schizophrenia and rats with abnormal prefrontal development. Mol Psychiatry. 2003;8(9):797–810. doi:10.1038/sj.mp.4001319

39. Eastwood SL, Cairns NJ, Harrison PJ. Synaptophysin gene expression in schizophrenia. Investigation of synaptic pathology in the cerebral cortex. Br J Psychiatry. 2000;176:236–242. doi:10.1192/bjp.176.3.236

40. Sekar A, Bialas AR, de Rivera H, et al. Schizophrenia risk from complex variation of complement component 4. Nature. 2016;530(7589):177–183. doi:10.1038/nature16549

41. Fromer M, Pocklington AJ, Kavanagh DH, et al. De novo mutations in schizophrenia implicate synaptic networks. Nature. 2014;506(7487):179–184. doi:10.1038/nature12929

42. Purcell SM, Moran JL, Fromer M, et al. A polygenic burden of rare disruptive mutations in schizophrenia. Nature. 2014;506(7487):185–190. doi:10.1038/nature12975

43. Mattheisen M, Mühleisen TW, Strohmaier J, et al. Genetic variation at the synaptic vesicle gene SV2A is associated with schizophrenia. Schizophr Res. 2012;141(2-3):262–265. doi:10.1016/j.schres.2012.08.027

44. Merritt K, Egerton A, Kempton MJ, Taylor MJ, McGuire PK. Nature of Glutamate Alterations in Schizophrenia: A Meta-analysis of Proton Magnetic Resonance Spectroscopy Studies. JAMA Psychiatry. 2016;73(7):665–674. doi:10.1001/jamapsychiatry.2016.0442

45. Nakahara T, Tsugawa S, Noda Y, et al. Glutamatergic and GABAergic metabolite levels in schizophrenia-spectrum disorders: a meta-analysis of 1H-magnetic resonance spectroscopy studies. Molecular Psychiatry. September 2021. doi:10.1038/s41380-021-01297-6

46. Stan AD, Ghose S, Zhao C, et al. Magnetic resonance spectroscopy and tissue protein concentrations together suggest lower glutamate signaling in dentate gyrus in schizophrenia. Molecular Psychiatry. 2015;20(4):433–439. doi:10.1038/mp.2014.54

47. Egerton A, Modinos G, Ferrera D, McGuire PK. Neuroimaging studies of GABA in schizophrenia: a systematic review with meta-analysis. Translational Psychiatry. 2017;7(6):e1147–e1147. doi:10.1038/tp.2017.124

48. Bale TL, Abel T, Akil H, et al. The critical importance of basic animal research for neuropsychiatric disorders. Neuropsychopharmacol. 2019;44(8):1349–1353. doi:10.1038/s41386-019-0405-9

49. Chakravarty MM, Guma E. Small animal imaging presents an opportunity for improving translational research in biological psychiatry. Journal of Psychiatry and Neuroscience. 2021;46(5):E579–E582. doi:10.1503/jpn.210172

50. Gilani AI, Chohan MO, Inan M, et al. Interneuron precursor transplants in adult hippocampus reverse psychosis-relevant features in a mouse model of hippocampal disinhibition. PNAS. 2014;111(20):7450–7455. doi:10.1073/pnas.1316488111

51. Silberberg G, Darvasi A, Pinkas-Kramarski R, Navon R. The involvement of ErbB4 with schizophrenia: Association and expression studies. American Journal of Medical Genetics Part B: Neuropsychiatric Genetics. 2006;141B(2):142–148. doi:https://doi.org/10.1002/ajmg.b.30275

52. Norton N, Moskvina V, Morris DW, et al. Evidence that interaction between neuregulin 1 and its receptor erbB4 increases susceptibility to schizophrenia. American Journal of Medical Genetics Part B: Neuropsychiatric Genetics. 2006;141B(1):96–101. doi:https://doi.org/10.1002/ajmg.b.30236

53. Fazzari P, Paternain AV, Valiente M, et al. Control of cortical GABA circuitry development by Nrg1 and ErbB4 signalling. Nature. 2010;464(7293):1376–1380. doi:10.1038/nature08928

54. Neddens J, Buonanno A. Selective populations of hippocampal interneurons express ErbB4 and their number and distribution is altered in ErbB4 knockout mice. Hippocampus. 2010;20(6):724–744. doi:10.1002/hipo.20675

55. Del Pino I, Garcia-Frigola C, Dehorter N, et al. Erbb4 deletion from fast-spiking interneurons causes schizophrenia-like phenotypes. Neuron. 2013;79(6):1152–1168. doi:10.1016/j.neuron.2013.07.010

56. Golub MS, Germann SL, Lloyd KCK. Behavioral characteristics of a nervous system-specific erbB4 knock-out mouse. Behav Brain Res. 2004;153(1):159–170. doi:10.1016/j.bbr.2003.11.010

57. Zhang C, Ni P, Liu Y, et al. GABAergic Abnormalities Associated with Sensorimotor Cortico-striatal Community Structural Deficits in ErbB4 Knockout Mice and First-Episode Treatment-Naive Patients with Schizophrenia. Neurosci Bull. 2020;36(2):97–109. doi:10.1007/s12264-019-00416-2

58. Skirzewski M, Karavanova I, Shamir A, et al. ErbB4 signaling in dopaminergic axonal projections increases extracellular dopamine levels and regulates spatial/working memory behaviors. Mol Psychiatry. 2018;23(11):2227–2237. doi:10.1038/mp.2017.132

59. Fogarty M, Grist M, Gelman D, Marín O, Pachnis V, Kessaris N. Spatial genetic patterning of the embryonic neuroepithelium generates GABAergic interneuron diversity in the adult cortex. J Neurosci. 2007;27(41):10935–10946. doi:10.1523/JNEUROSCI.1629-07.2007

60. Grandjean J, Schroeter A, Batata I, Rudin M. Optimization of anesthesia protocol for resting-state fMRI in mice based on differential effects of anesthetics on functional connectivity patterns. NeuroImage. 2014;102:838–847. doi:10.1016/j.neuroimage.2014.08.043

61. Grandjean J, Canella C, Anckaerts C, et al. Common functional networks in the mouse brain revealed by multi-centre resting-state fMRI analysis. NeuroImage. 2020;205:116278. doi:10.1016/j.neuroimage.2019.116278

62. Hirschler L, Debacker CS, Voiron J, Köhler S, Warnking JM, Barbier EL. Interpulse phase corrections for unbalanced pseudo-continuous arterial spin labeling at high magnetic field. Magnetic Resonance in Medicine. 2018;79(3):1314–1324. doi:10.1002/mrm.26767

63. Andersson JLR, Skare S, Ashburner J. How to correct susceptibility distortions in spin-echo echo-planar images: application to diffusion tensor imaging. Neuroimage. 2003;20(2):870–888. doi:10.1016/S1053-8119(03)00336-7

64. Wood TC. QUIT: QUantitative Imaging Tools. Journal of Open Source Software. 2018;3(26):656. doi:10.21105/joss.00656

65. Avants BB, Epstein CL, Grossman M, Gee JC. Symmetric diffeomorphic image registration with cross-correlation: Evaluating automated labeling of elderly and neurodegenerative brain. Medical Image Analysis. 2008;12(1):26–41. doi:10.1016/j.media.2007.06.004

66. Vernon AC, So PW, Lythgoe DJ, et al. Longitudinal in vivo maturational changes of metabolites in the prefrontal cortex of rats exposed to polyinosinic-polycytidylic acid in utero. Eur Neuropsychopharmacol. 2015;25(12):2210–2220. doi:10.1016/j.euroneuro.2015.09.022

67. Yahya A. Metabolite detection by proton magnetic resonance spectroscopy using PRESS. Progress in Nuclear Magnetic Resonance Spectroscopy. 2009;55(3):183–198. doi:https://doi.org/10.1016/j.pnmrs.2009.04.001

68. Simpson R, Devenyi GA, Jezzard P, Hennessy TJ, Near J. Advanced processing and simulation of MRS data using the FID appliance (FID-A)-An open source, MATLAB-based toolkit. Magn Reson Med. 2017;77(1):23–33. doi:10.1002/mrm.26091

69. Provencher SW. Estimation of metabolite concentrations from localized in vivo proton NMR spectra. Magnetic Resonance in Medicine. 1993;30(6):672–679. doi:https://doi.org/10.1002/mrm.1910300604

70. Provencher SW. Automatic quantitation of localized in vivo 1H spectra with LCModel. NMR in Biomedicine. 2001;14(4):260–264. doi:https://doi.org/10.1002/nbm.698

71. Muñoz-Hernández MC, García-Martín ML. In Vivo 1H Magnetic Resonance Spectroscopy. In: García Martín ML, López Larrubia P, eds. Preclinical MRI: Methods and Protocols. New York, NY: Springer New York; 2018:151–167. doi:10.1007/978-1-4939-7531-0_10

72. Jansen D, Zerbi V, Janssen CI, et al. A longitudinal study of cognition, proton MR spectroscopy and synaptic and neuronal pathology in aging wild-type and AβPPswe-PS1dE9 mice. PLoS One. 2013;8(5):e63643. doi:10.1371/journal.pone.0063643

73. Sidek S, Ramli N, Rahmat K, Ramli NM, Abdulrahman F, Kuo TL. In vivo proton magnetic resonance spectroscopy (1H-MRS) evaluation of the metabolite concentration of optic radiation in primary open angle glaucoma. Eur Radiol. 2016;26(12):4404–4412. doi:10.1007/s00330-016-4279-5

74. Peris-Yague A, Kiemes A, Cash D, et al. Region-specific and dose-specific effects of chronic haloperidol exposure on [3H]-flumazenil and [3H]-Ro15-4513 GABAA receptor binding sites in the rat brain. European Neuropsychopharmacology. 2020;41:106–117. doi:https://doi.org/10.1016/j.euroneuro.2020.10.004

75. Kiemes A, Gomes FV, Cash D, et al. GABAA and NMDA receptor density alterations and their behavioral correlates in the gestational methylazoxymethanol acetate model for schizophrenia. Neuropsychopharmacology. 2022;47:687–695. doi:10.1038/s41386-021-01213-0

76. Myers JF, Comley RA, Gunn RN. Quantification of [11C]Ro15-4513 GABAAα5 specific binding and regional selectivity in humans. Journal of Cerebral Blood Flow & Metabolism. 2017;37(6):2137–2148. doi:10.1177/0271678×16661339

77. Myers JF, Rosso L, Watson BJ, et al. Characterisation of the Contribution of the GABA-Benzodiazepine α1 Receptor Subtype to [11C]Ro15-4513 PET Images. Journal of Cerebral Blood Flow & Metabolism. 2012;32(4):731–744. doi:10.1038/jcbfm.2011.177

78. Lukow PB, Martins D, Veronese M, et al. Cellular and molecular signatures of in vivo GABAergic neurotransmission in the human brain. bioRxiv. 2021:2021.06.17.448812. doi:10.1101/2021.06.17.448812

79. Sieghart W. Structure, pharmacology, and function of GABAA receptor subtypes. Adv Pharmacol. 2006;54:231–263. doi:10.1016/s1054-3589(06)54010-4

80. Nabulsi NB, Mercier J, Holden D, et al. Synthesis and Preclinical Evaluation of 11C-UCB-J as a PET Tracer for Imaging the Synaptic Vesicle Glycoprotein 2A in the Brain. Journal of Nuclear Medicine. 2016;57(5):777–784. doi:10.2967/jnumed.115.168179

81. Lieberman JA, Girgis RR, Brucato G, et al. Hippocampal dysfunction in the pathophysiology of schizophrenia: a selective review and hypothesis for early detection and intervention. Molecular Psychiatry. 2018;23(8):1764–1772. doi:10.1038/mp.2017.249

82. Gill KM, Grace AA. Corresponding decrease in neuronal markers signals progressive parvalbumin neuron loss in MAM schizophrenia model. International Journal of Neuropsychopharmacology. 2014;17(1609-1619). doi:10.1017/S146114571400056X

83. Ben-Shachar M, Lüdecke D, Makowski D. effectsize: Estimation of Effect Size Indices and Standardized Parameters. Journal of Open Source Software. 2020;5(56). doi:10.21105/joss.02815

84. Paxinos G, Franklin KB. Paxinos and Franklin’s the Mouse Brain in Stereotaxic Coordinates. Academic press; 2019.

85. Kuschinsky W. Coupling of function, metabolism, and blood flow in the brain. Neurosurg Rev. 1991;14(3):163–168. doi:10.1007/BF00310651

86. Wen L, Lu YS, Zhu XH, et al. Neuregulin 1 regulates pyramidal neuron activity via ErbB4 in parvalbumin-positive interneurons. PNAS. 2010;107(3):1211–1216. doi:10.1073/pnas.0910302107

87. Stone JM, Dietrich C, Edden R, et al. Ketamine effects on brain GABA and glutamate levels with 1H-MRS: relationship to ketamine-induced psychopathology. Mol Psychiatry. 2012;17(7):664–665. doi:10.1038/mp.2011.171

88. Homayoun H, Moghaddam B. NMDA Receptor Hypofunction Produces Opposite Effects on Prefrontal Cortex Interneurons and Pyramidal Neurons. J Neurosci. 2007;27(43):11496–11500. doi:10.1523/JNEUROSCI.2213-07.2007

89. Kim SY, Lee H, Kim HJ, et al. In vivo and ex vivo evidence for ketamine-induced hyperglutamatergic activity in the cerebral cortex of the rat: Potential relevance to schizophrenia. NMR Biomed. 2011;24(10):1235–1242. doi:10.1002/nbm.1681

90. Rothman DL, Sibson NR, Hyder F, Shen J, Behar KL, Shulman RG. In vivo nuclear magnetic resonance spectroscopy studies of the relationship between the glutamate-glutamine neurotransmitter cycle and functional neuroenergetics. Philos Trans R Soc Lond B Biol Sci. 1999;354(1387):1165–1177. doi:10.1098/rstb.1999.0472

91. Rothman DL, Behar KL, Hyder F, Shulman RG. In vivo NMR Studies of the Glutamate Neurotransmitter Flux and Neuroenergetics: Implications for Brain Function. Annual Review of Physiology. 2003;65(1):401–427. doi:10.1146/annurev.physiol.65.092101.142131

92. Snyder J, Wilman A. Field strength dependence of PRESS timings for simultaneous detection of glutamate and glutamine from 1.5 to 7T. Journal of Magnetic Resonance. 2010;203(1):66–72. doi:https://doi.org/10.1016/j.jmr.2009.12.002

93. Inda MC, DeFelipe J, Muñoz A. Morphology and Distribution of Chandelier Cell Axon Terminals in the Mouse Cerebral Cortex and Claustroamygdaloid Complex. Cerebral Cortex. 2009;19(1):41–54. doi:10.1093/cercor/bhn057

94. Hill RA. Sex differences in animal models of schizophrenia shed light on the underlying pathophysiology. Neurosci Biobehav Rev. 2016;67:41–56. doi:10.1016/j.neubiorev.2015.10.014

95. Goldstein JM, Cherkerzian S, Tsuang MT, Petryshen TL. Sex differences in the genetic risk for schizophrenia: history of the evidence for sex-specific and sex-dependent effects. Am J Med Genet B Neuropsychiatr Genet. 2013;162B(7):698–710. doi:10.1002/ajmg.b.32159

96. Arrive Guidelines. IMPC | International Mouse Phenotyping Consortium. https://www.mousephenotype.org/about-impc/animal-welfare/arrive-guidelines/. Accessed March 4, 2022.

97. Arnegard ME, Whitten LA, Hunter C, Clayton JA. Sex as a Biological Variable: A 5-Year Progress Report and Call to Action. Journal of Women’s Health. 2020;29(6):858–864. doi:10.1089/jwh.2019.8247

98. NC3Rs. https://www.nc3rs.org.uk/. Accessed March 4, 2022.

99. Karp NA, Reavey N. Sex bias in preclinical research and an exploration of how to change the status quo. British Journal of Pharmacology. 2019;176(21):4107–4118. doi:10.1111/bph.14539

100. Young JW, Henry BL, Geyer MA. Animal models of schizophrenia. British Journal of Pharmacology. 2011;164:1162–1194. doi:10.1111/bph.2011.164.issue-4

