## Supplementary Information for "*Erbb4* deletion from fast-spiking interneurons causes psychosis-relevant neuroimaging phenotypes"

**Supplementary Methods**

For completeness, in addition to the hippocampus, exploratory analyses were also performed on the amygdala, retrosplenial cortex (RSPcx), visual cortex (Sencx), prelimbic cortex (PLcx), motor cortex (Mcx), and orbital cortex (ORBcx) (Figure S1). Means and standard deviations along with the results of exploratory independent t-tests run in GraphPad Prism software (v9.2.0 for Windows) are presented in tables S3-5 below. Due to issues with the tissue preparation, data are missing for ROIs in some mice. Where the number of datapoints per genotype group per ROI fell below n=5, data was not reported.

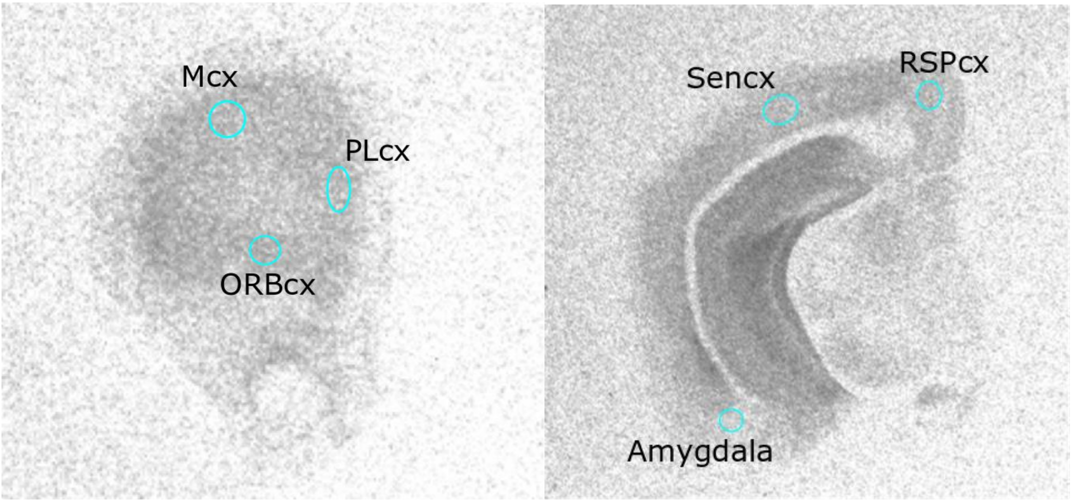


*Figure S1. Representative autoradiograph with additional regions of interest. Frontal regions (left): motor cortex (Mcx), prelimbic cortex (PLcx), orbital cortex (ORBcx). Posterior regions (right): retrosplenial cortex (RSPcx), visual cortex (Sencx), amygdala.*

### Supplementary Results

*Table S1. Cerebral blood flow of whole brain and additional atlas ROIs*

| | *ErbB4*$^{F/F}$ | *Lhx6-Cre;ErbB4*$^{F/F}$ | *ErbB4*$^{F/F}$ vs. *Lhx6-Cre;ErbB4*$^{F/F}$ | |
|---|---|---|---|---|
| | Mean (SD) | Mean (SD) | *t* | *p* |
| Whole brain | 62.41 (19.89) | 59.68 (11.62) | 0.40 | 0.69 |
| Amygdala | 0.90 (0.07) | 0.94 (0.09) | 0.98 | 0.34 |
| Brain stem | 0.95 (0.16) | 0.87 (0.28) | 0.82 | 0.42 |
| Cerebellum | 1.13 (0.22) | 1.02 (0.37) | 1.22 | 0.24 |
| Cingulate cx | 1.10 (0.25) | 1.14 (0.19) | 0.40 | 0.69 |
| Colliculus | 1.18 (0.12) | 1.29 (0.22) | 1.32 | 0.20 |
| Entorhinal cx | 1.15 (0.13) | 1.29 (0.14) | 2.40 | 0.03* |
| Hypothalamus | 0.66 (0.06) | 0.67 (0.05) | 0.26 | 0.80 |
| Midbrain | 0.87 (0.07) | 0.83 (0.11) | 1.01 | 0.32 |
| Motor cx | 0.96 (0.14) | 1.00 (0.16) | 0.60 | 0.56 |
| Olfactory | 1.13 (0.09) | 1.19 (0.13) | 1.23 | 0.23 |
| PAG | 0.76 (0.11) | 0.74 (0.13) | 0.28 | 0.78 |
| Pallidum | 0.92 (0.05) | 0.95 (0.08) | 0.92 | 0.37 |
| Prefrontal cx | 1.05 (0.16) | 1.11 (0.12) | 0.98 | 0.34 |
| Sensory cx | 1.08 (0.13) | 1.16 (0.14) | 1.39 | 0.18 |
| Septum | 0.60 (0.15) | 0.62 (0.12) | 0.46 | 0.65 |
| Striatum | 0.92 (0.09) | 0.95 (0.09) | 0.57 | 0.57 |
| Thalamus | 0.87 (0.08) | 0.92 (0.14) | 1.13 | 0.27 |

ROI: region of interest; cx: cortex; PAG: periaqueductal gray; *$p<0.05$

*Table S2. $^{1}$H-MRS quality control parameters*

| | *ErbB4$^{F/F}$* | *Lhx6-Cre;ErbB4$^{F/F}$* | *ErbB4$^{F/F}$* vs. *Lhx6-Cre;ErbB4$^{F/F}$* | |
|---|---|---|---|---|
| | Mean (SD) | Mean (SD) | *t* | *p* |
| SNR | 14.7 (1.7) | 15.8 (3.8) | 0.87 | 0.39 |
| FWHM (ppm) | 0.05 (0.01) | 0.05 (0.03) | 0.60 | 0.55 |
| GABA CRLB (%) | 10.58 (2.39) | 10.92 (2.94) | 0.31 | 0.76 |
| Gln CRLB (%) | 9.50 (1.73) | 8.75 (4.07) | 0.59 | 0.01* |
| Glu CRLB (%) | 4.08 (0.67) | 4.00 (1.13) | 0.22 | 0.10 |

SNR: signal-to-noise ratio; FWHM: full width half maximum; CRLB: Cramér-Rao lower bound; Gln: glutamine; *$p<0.05$

*Table S3. Exploratory independent t-tests of additional [$^3$H]Ro15-4513 ROIs*

| | *ErbB4*$^{F/F}$ | *Lhx6-Cre;ErbB4*$^{F/F}$ | *ErbB4*$^{F/F}$ vs. *Lhx6-Cre;ErbB4*$^{F/F}$ | |
|---|---|---|---|---|
| | Mean (SD) | Mean (SD) | *t* | *p* |
| Amygdala | 3.07 (0.95) | 3.11 (1.15) | 0.09 | 0.93 |
| RSPcx | 2.91 (1.08) | 3.91 (0.86) | 2.33 | 0.03* |
| Sencx | 5.7 (1.19) | 5.94 (1.56) | 0.39 | 0.69 |
| PLcx | 7.27 (1.89) | 6.1 (1.97) | 1.35 | 0.19 |
| Mcx | 5.54 (1.28) | 6.37 (1.18) | 1.45 | 0.16 |
| ORBcx | 3.88 (2.1) | 4.98 (1.37) | 1.35 | 0.19 |

ROI: region of interest; RSPcx: retrosplenial cortex; Sencx: visual cortex; PLcx: prelimbic cortex; Mcx: motor cortex; ORBcx: orbitofrontal cortex; *$p<0.05$

*Table S4. Exploratory independent t-tests of additional [$^3$H]flumazenil ROIs*

| | *ErbB4$^{F/F}$* | *Lhx6-Cre;ErbB4$^{F/F}$* | *ErbB4$^{F/F}$* vs. *Lhx6-Cre;ErbB4$^{F/F}$* | |
|---|---|---|---|---|
| | Mean (SD) | Mean (SD) | *t* | *p* |
| Amygdala | 5.19 (2.62) | 7.23 (4.6) | 1.02 | 0.32 |
| RSPcx | 10.75 (4.05) | 7.87 (1.68) | 1.99 | 0.06 |
| Sencx | 15.97 (4.2) | 12.75 (2.05) | 1.97 | 0.07 |
| PLcx | 13.01 (3.75) | 12.36 (3.55) | 0.36 | 0.72 |
| Mcx | 14.19 (3.4) | 13.11 (2.82) | 0.73 | 0.47 |
| ORBcx | 12.96 (5.07) | 11.9 (3.77) | 0.46 | 0.65 |

ROI: region of interest; RSPcx: retrosplenial cortex; Sencx: visual cortex; PLcx: prelimbic cortex; Mcx: motor cortex; ORBcx: orbitofrontal cortex

*Table S5. Exploratory independent t-tests of all [$^3$H]UCB-J ROIs*

| | *ErbB4$^{F/F}$* | *Lhx6-Cre;ErbB4$^{F/F}$* | *ErbB4$^{F/F}$* vs. *Lhx6-Cre;ErbB4$^{F/F}$* | |
|---|---|---|---|---|
| | Mean (SD) | Mean (SD) | *t* | *p* |
| RSPcx | 192.09 (14.95) | 176.92 (20.94) | 1.80 | 0.09 |
| Sencx | 197.47 (14.00) | 189.69 (15.12) | 1.18 | 0.25 |
| PLcx | 195.91 (11.97) | 193.83 (20.44) | 0.24 | 0.82 |
| Mcx | 191.07 (15.86) | 181.08 (12.27) | 1.54 | 0.14 |
| ORBcx | 192.52 (13.61) | 179.93 (21.48) | 1.31 | 0.21 |

ROI: region of interest; RSPcx: retrosplenial cortex; Sencx: visual cortex; PLcx: prelimbic cortex; Mcx: motor cortex; ORBcx: orbitofrontal cortex

**Supplementary Discussion**

Exploratory analysis of CBF values in additional parcellated ROIs revealed a significant increase in CBF in the entorhinal cortex of *Erbb4* mutants. Although no brain perfusion studies have implicated the entorhinal cortex in psychosis, multiple lines of evidence converge to implicate this area in relation to interneurons and cognitive deficits. Preclinical evidence indicates that entorhinal cortex communication with the hippocampus is integral for gamma oscillations[1], prepulse inhibition is dependent on this region[2], and a genetic animal model (LPA1-deficient mice) shows reduction of entorhinal cortex PV+ interneurons[3]. Further, putative compensatory increases in α1-3;5GABA$_A$R are linked with cognitive impairments in psychosis patients[4]. Our findings demonstrate that inhibitory interneuron disruption also increases network activity in the entorhinal cortex in psychosis in *Erbb4* mutant mice.

Interestingly our exploratory analysis revealed α5GABA$_A$R increases (uncorrected) in the retrosplenial cortex and decreases in α1-3;5GABA$_A$R that approached significance in the same region. [$^3$H]Flumazenil and [$^3$H]Ro15-4513 binding distributions seem to be anticorrelated[5] potentially yielding such differential results in the same region. The retrosplenial cortex features reciprocal links to the hippocampus, parahippocampus and thalamus and is involved in an array of cognitive abilities including spatial navigation, memory, and planning[6]. While neuroimaging research has identified functional connectivity changes in this region in psychosis patients suggesting aberrant activity in this region[7–10], GABAergic receptors have not been studied in this brain region. Nonetheless, preclinical work has suggested that the retrosplenial cortex may suffer a loss of PV+ interneurons in relation to psychosis[11], and we extend such findings to show PV+ interneuron impairment generates GABAergic receptor differences that may contribute to aberrant neural activity.

**Supplementary References**


1. Fernández-Ruiz A, Oliva A, Nagy GA, Maurer AP, Berényi A, Buzsáki G. Entorhinal-CA3 Dual-Input Control of Spike Timing in the Hippocampus by Theta-Gamma Coupling. *Neuron*. 2017;93(5):1213-1226.e5. doi:10.1016/j.neuron.2017.02.017

2. Goto K, Ueki A, Iso H, Morita Y. Reduced prepulse inhibition in rats with entorhinal cortex lesions. *Behav Brain Res*. 2002;134(1-2):201-207. doi:10.1016/s0166-4328(02)00039-6

3. Cunningham MO, Hunt J, Middleton S, et al. Region-Specific Reduction in Entorhinal Gamma Oscillations and Parvalbumin-Immunoreactive Neurons in Animal Models of Psychiatric Illness. *J Neurosci*. 2006;26(10):2767-2776. doi:10.1523/JNEUROSCI.5054-05.2006

4. Frankle WG, Cho RY, Prasad KM, et al. In vivo measurement of GABA transmission in healthy subjects and schizophrenia patients. *Am J Psychiatry*. 2015;172(11):1148-1159. doi:10.1176/appi.ajp.2015.14081031

5. Lingford-Hughes A, Hume SP, Feeney A, et al. Imaging the GABA-Benzodiazepine Receptor Subtype Containing the α5-Subunit In Vivo with [11C]Ro15 4513 Positron Emission Tomography. *J Cereb Blood Flow Metab*. 2002;22(7):878-889. doi:10.1097/00004647-200207000-00013

6. Vann SD, Aggleton JP, Maguire EA. What does the retrosplenial cortex do? *Nat Rev Neurosci*. 2009;10(11):792-802. doi:10.1038/nrn2733

7. Tendolkar I, Weis S, Guddat O, et al. Evidence for a dysfunctional retrosplenial cortex in patients with schizophrenia: a functional magnetic resonance imaging study with a semantic—perceptual contrast. *Neuroscience Letters*. 2004;369(1):4-8. doi:10.1016/j.neulet.2004.07.024

8. Siemerkus J, Irle E, Schmidt-Samoa C, Dechent P, Weniger G. Egocentric spatial learning in schizophrenia investigated with functional magnetic resonance imaging. *NeuroImage: Clinical*. 2012;1(1):153-163. doi:10.1016/j.nicl.2012.10.004

9. Wang Y, Yan C, Yin D zhi, et al. Neurobiological changes of schizotypy: evidence from both volume-based morphometric analysis and resting-state functional connectivity. *Schizophr Bull*. 2015;41 Suppl 2:S444-454. doi:10.1093/schbul/sbu178

10. Bluhm RL, Miller J, Lanius RA, et al. Retrosplenial cortex connectivity in schizophrenia. *Psychiatry Research: Neuroimaging*. 2009;174(1):17-23. doi:10.1016/j.pscychresns.2009.03.010

11. Klimczak P, Rizzo A, Castillo-Gómez E, et al. Parvalbumin Interneurons and Perineuronal Nets in the Hippocampus and Retrosplenial Cortex of Adult Male Mice After Early Social Isolation Stress and Perinatal NMDA Receptor Antagonist Treatment. *Frontiers in Synaptic Neuroscience*. 2021;13. https://www.frontiersin.org/article/10.3389/fnsyn.2021.733989. Accessed March 4, 2022.
